# Towards Optimizing Target Engagement in Non-Invasive Trigeminal Nerve Stimulation: Anatomical Characterization and Computational Modeling of the Human Trigeminal Nerve

**DOI:** 10.1101/2025.06.19.660375

**Authors:** Jennifer L. Perrault, Keith D. Kozma, Weifeng Zeng, Zeeda Nkana, Nicholas J. Albano, Kirsten A. Gunderson, Samuel A. Hurley, Wendell B. Lake, Justin C. Williams, Samuel O. Poore, Kip A. Ludwig, Aaron M. Dingle, Aaron J. Suminski

## Abstract

**Objective:** Cranial nerve stimulation (CNS) uses electric current to modulate higher-order brain activity and organ function via nerves, including the vagus and trigeminal, with applications in migraine, epilepsy, and pediatric ADHD. The trigeminal nerve is an emerging target for non-invasive neuromodulation due to the superficial trajectory of its branches, the supraorbital (SON), infraorbital (ION), and mental nerves (MN), and the predominantly sensory composition of the SON and ION. However, the parameters and outcomes of trigeminal nerve stimulation (TNS) remain varied.

**Approach:** This study characterizes the anatomical course, tissue composition, and activation profiles of the SON, ION, and MN using five human donors. CT imaging was utilized to localize each nerve’s exit foramen and distance to midline. Microdissections quantified nerve circumference and depth relative to the skin surface. Histological analysis described the number of fascicles and fascicular tissue area. Nerve depths were incorporated into computational models to illustrate the activation function across tissue layers, comparing expected nociceptor and nerve trunk activation functions as a measure of neural engagement.

**Main Results:** The SON was found to be significantly more superficial than the ION and MN and had a higher nerve-to-connective tissue ratio relative to the MN. Computational modeling demonstrated that the activation function at the depths of nociceptors was orders of magnitude greater than within the main nerve trunks, suggesting preferential recruitment of cutaneous nociceptors, dependent on nociceptor density.

**Significance:** The SON presents the most accessible and anatomically favorable target for transcutaneous trigeminal nerve stimulation among the branches examined due to its superficial location. However, preferential activation of low-threshold nociceptors compared to nerve trunks may lead to treatment-limiting off-target side effects, favoring strategies that target fibers of interest within the skin. These findings offer an anatomically informed framework to guide further computational modeling and electrode design for targeted trigeminal neuromodulation.

## Introduction

Cranial nerve stimulation (CNS) uses electrical current to activate cranial nerves (i.e., the trigeminal, hypoglossal, and vagus nerves) in an attempt to treat neurological disorders by modulating connected brain regions and organ systems. This field has evolved over the past several decades to include FDA-approved therapies for a wide variety of disorders including epilepsy (Morris, 1999; Ben-Menachem, 2002; Englot, 2011), obstructive sleep apnea (OSA) (Schwartz, 2001; Eastwood, 2011), and treatment-resistant depression (TRD) (Rush, 2000; Nemeroff, 2006; Conway, 2020). Furthermore, trigeminal nerve stimulation (TNS), a rapidly growing segment of CNS, includes FDA-approved, FDA-cleared, and CE mark-certified devices for indications such as migraine (Riederer, 2015; Chou, 2017; Chou 2019), pediatric attention deficit hyperactivity disorder (ADHD) (McGough, 2019; Loo, 2021), and epilepsy in the European market (DeGiorgio, 2013; Cook, 2015; Olivié, 2019; Gil-López, 2020). In addition to these current treatments, TNS is being investigated for promoting cortical plasticity for recovery after neural injury (Suminski, 2018) and modulating cerebral blood flow (Lang and Zimmer, 1974; Atalay, 2002; Li, 2021; White, 2021), which is thought to impact recovery after traumatic brain injury (Chiluwal, 2017; Li, 2021; Yang, 2022, Xu, 2023). However, in many clinical applications of trigeminal nerve stimulation (TNS), particularly transcutaneous electrical nerve stimulation (TENS), stimulation intensity and outcomes vary significantly across studies and indications (Shiozawa, 2014; Powell, 2023; Westwood, 2023). The basis of inconsistent target engagement in applications using transcutaneous TNS is multifactorial and likely driven by person-to-person anatomical variability in the composition, depths, and branching patterns of the trigeminal nerves. This underscores the need to better understand the factors contributing to therapeutic stimulation, with the goal of maximizing on-target effects while minimizing side effects.

Consisting of three main divisions, the ophthalmic (V1), maxillary (V2), and mandibular (V3), the trigeminal nerve is an especially attractive target for neuromodulation due to several key anatomical and physiological characteristics. First, the V1, V2, and V3 divisions give rise to multiple superficial branches, including the supraorbital nerve (SON), infraorbital nerve (ION), and mental nerve (MN), respectively, making them more accessible for non-invasive transcutaneous stimulation. Second, the SON and ION branches contain purely sensory axons (Bathla, 2013; Joo, 2013; Powell, 2023) and have superficial exit points from the skull (Beer, 1998; Joo, 2013; Ference, 2015; Netter, 2019), decreasing the likelihood of off-target muscle activation as seen in vagus nerve stimulation from motor fiber activation (Nicolai, 2020; Settell, 2020). Finally, the trigeminal nerve has broad access to various neurotransmitter systems and neural pathways via its connection to brainstem nuclei. For example, afferent fibers from the trigeminal nerve branches project to key central structures, notably the sphenopalatine ganglion (SPG), trigeminal subnucleus caudalis (TSN), and an indirect projection to the nucleus tractus solitarius (NTS) via the trigeminal nuclei (Fanselow, 2012; Huff, 2025; Patel, 2025). The NTS is a major integrator of autonomic and sensory input, providing a mechanism to influence autonomic tone by affecting blood pressure and respiration, whereas the SPG and TSN modulate cerebral blood flow directly (Lim, 2023), with emerging clinical applications in traumatic brain injury recovery (Chiluwal, 2017). Importantly, these direct and indirect projections are similar to those of the vagus nerve, making the trigeminal nerve a potential alternative target for some indications (i.e. epilepsy and TRD) (Fanselow, 2012).

Unfortunately, current clinical applications of transcutaneous trigeminal nerve stimulation are highly variable in stimulation location and intensity (Shiozawa, 2014; Powell, 2023; Westwood, 2023), and success criteria for target engagement have not been thoroughly studied. Thus, there are several key barriers to selective target engagement in TNS that have been inadequately addressed. First, activation of on-target neural fibers is governed by the underlying biophysics that dictate that neural engagement drops off rapidly with distance, following the inverse square law for bipolar sources (Plonsey and Barr, 2007). Consequently, fibers closest to the electrode—including nociceptors and Aβ/Aδ afferent fibers innervating the skin between the stimulation electrode and the intended nerve—are activated first. Second, reverse recruitment order of axons mandating that larger-diameter fibers are depolarized at lower thresholds than smaller-diameter fibers, regardless of their functional type (Plonsey and Barr, 2007; Grinberg, 2008), may lead to off-target stimulation. In TNS, trigeminal afferents such as Aβ and Aδ fibers are presumed therapeutic targets, but cutaneous nociceptors between the electrode and the trigeminal nerve trunk and/or large-diameter Aα motor fibers from other nerve trunks (i.e. facial nerve) may be inadvertently activated first, limiting tolerability and therapeutic efficacy. Finally, fascicle size—impacting perineurium thickness within nerve trunks—changes the threshold stimulus current for activation (Davis, 2023; Grinberg, 2008), increasing the zone of activation and potential off-target effects. Precise anatomical mapping and investigation of target nerve activation characteristics are needed to decrease off-target effects and deliver precise neuromodulation to the trigeminal nerve.

To provide a framework for refining target engagement strategies in transcutaneous TNS, this study maps the major cutaneous trigeminal nerve branches, highlighting anatomical features as well as relevant parameters for the modeling of trigeminal nerve engagement in neuromodulation. We quantify nerve depth from the skin surface at the well-defined foraminal landmarks, the associated foramina-midline distance, and fascicular cross-sectional area and organization. Furthermore, we examined the branching morphology of each nerve immediately after exiting the foramen via qualitative microdissection in five human donors. We then used our anatomically derived depth measurements to develop a COMSOL finite element model (FEM) to simulate the activation of the SON, ION, and MN, as well as surrounding nociceptors, using the activation function as a proxy for fiber engagement at relevant depths. Lastly, we conducted a sensitivity analysis to understand how skin nociceptors and trigeminal nerve fibers are activated based on variability in skin impedance parameters. This work provides a foundational step toward the design of further computational models, electrode configurations, and stimulation parameters that maximize therapeutic engagement in TNS.

## Methods

### Anatomical Characterization of Trigeminal Nerve

CT Imaging, microdissection and tissue harvest, and histological examination were performed on five human donors obtained from Science Care (Phoenix, AZ). All donors were male with the following demographics: age - 69.4± 20.5 years, weight - 178.2±24.6 lbs., and BMI - 25.38±2.95. (Avg ± Standard Deviation). Prior to surgical dissection and tissue harvest, high-resolution computed tomography (CT) was performed on each donor (see CT Imaging and analysis for details). Next, surgical dissection was performed to isolate the SON, ION, and MN branches of the trigeminal nerve (see below for details). Following dissection, the depth of each nerve trunk was measured by inserting a needle through the skin to the foramen, a mark was made on the needle where the surface of the skin met the needle, and the needle was then removed and measured from the tip to the mark. After each nerve depth was measured, nerve tissue samples were harvested at the point where the nerve exited the foramen, and before the nerve began to spread and branch.

#### SON Surgical Dissection

The SON was accessed through a blepharoplasty-type incision, immediately inferior to the eyebrow, providing a direct route to the supraorbital region. Blunt dissection was carried out in the subcutaneous tissue toward the superior orbital rim to identify the supraorbital foramen, or notch. Once the bony edge was reached and the nerve was visualized exiting the foramen, dissection was continued into the orbit, deep to the orbital fat, to delineate the nerve along its course within the orbital roof.

#### ION Surgical Dissection

The ION was exposed through a transconjunctival incision along the inner aspect of the lower eyelid, providing direct access to the infraorbital region. The dissection was carried downward toward the lower orbital rim, carefully following the periosteal plane. The dissection continued until the infraorbital foramen was identified along the lower margin of the orbit, with the ION emerging from the foramen.

#### MN Surgical Dissection

The MN was accessed with a combined midline and transmucosal incision inside the oral cavity. A midline incision was made along the alveolar mucosa at the junction between the cheek mucosa and the gum line. Dissection was carefully performed along the periosteal layer of the mandible, toward the submandibular region, until the mental foramen was identified on the anterior mandibular bone with the MN emerging from the foramen.

### Histology

All nerve tissue samples were fixed with 10% neutral buffered formalin at 4°C overnight. Nerves were embedded in paraffin with the rostral face of the nerve down, with transverse 5µm serial sections taken rostral to caudal. Sections were stained with Gomori’s Trichrome, imaged using the Aperio ImageScope (Leica Biosystems, Deer Park, IL, USA), and analyzed with the associated Aperio ImageScope software (v12.3.3.5048) for nerve circumference, total area, and fascicular area. Here, the circumference of the nerve was manually traced using the annotation tool, which provided the nerve’s circumference and total area. Next, each fascicle was counted, manually traced using the same method described above, and the area was recorded. Lastly, we used the ruler function to measure the length and width of each nerve sample. All annotations for representative nerves from each branch are shown in **Supplementary Figure 1**. The fascicular tissue ratio, describing the percent makeup of “within fascicle” tissue to the total area, was computed as: *fascicular tissue area (mm^2^) / total area (mm^2^).* Additionally, fascicle counts and mean fascicle areas were reported in **Supplemental Table 1**, which have implications in thresholds for activation; larger fascicle diameters correlate with higher activation thresholds (Davis, 2023; Grinberg, 2008) and are ultimately dependent on the location of nerve cross sections.

### CT Imaging and Analysis

Prior to dissections, we acquired CT scans of each donor in order to localize the foramen where each nerve branch exited the skull. Measurements of the trigeminal nerve foramina relative to the midline provide an anatomical landmark as each trigeminal nerve branch exits from the skull. This guides transcutaneous electrode placement targeting the branching patterns of each nerve. Here, frozen donor tissue was placed in a GE HealthCare Discovery CT750 HD (Chicago, IL, USA) and images were acquired with a 120 kVp, 270 mA, helical acquisition with 0.53 pitch factor, acquired diameter 320 mm, reconstructed diameter 250 mm, 512 x 512 matrix, 5 mm slice thickness. ITK-SNAP (Yushkevich, 2006) was used to analyze CT images of each donor and measure the distance of the foramina to the midline. The midline for the supraorbital and infraorbital foramina was determined using the sagittal suture, while the space between the most medial lower incisors was used as the midline for the mental nerve foramen. Approximate locations for each foramen were estimated (Netter, 2019), and then sagittal CT views were used to find a small, dark grey indentation in the bone, indicative of the foramen. After isolating the foramen on the sagittal view, coronal views were used to confirm the precise location of the foramen. After verifying the location of the foramen in each plane, the measuring tool on ITK-SNAP was used to measure from the foramen to the appropriate midline marker. Representative examples of the foramina where the SON, ION, and MN leave the skull are shown in **Figure 1**.

**Figure 1.**
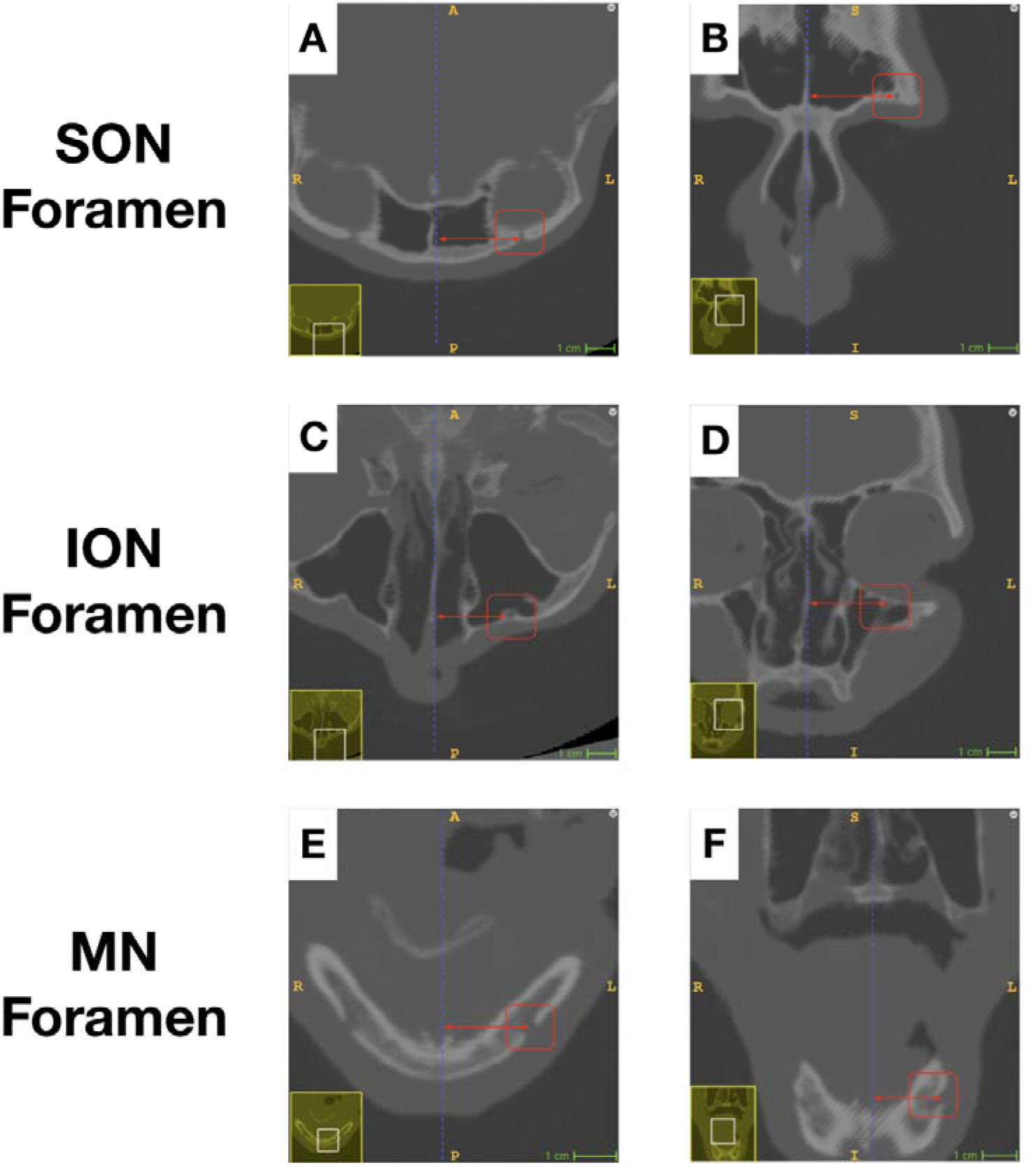
Representative CT image of one donor’s (D1) left supraorbital nerve (SON), left infraorbital nerve (ION), and left mental nerve (MN) foramina within the skull, showing methods of measuring foramina-midline distances. Two views are shown: axial in the first column and coronal in the second column. Panel A shows an axial view of the left SON foramen, directly above the eye socket, circled in red. Red arrows indicate the distance reported from the midline structures (blue dotted line). Panel B shows the corresponding coronal view of the SON foramen to midline structures near the nasal cavity. Yellow and white boxes in the lower left of each panel indicate the corresponding zoomed-out view for coronal and axial slices. Panels C and D show axial and coronal views of the left ION foramen relative to bony landmarks on the midline under the eye socket. Panels E and F show the left MN foramen in the mandible. <u>Supplemental Video 1</u> dynamically zooms through dorsal-to-ventral coronal planes of the SON and ION, and rostral-to-caudal axial planes of the MN, where the hole in the bony structure (light) can be seen clearly where each of the three branches exits the skull through their respective foramen.

### Statistical Testing

All statistical analyses were performed in R. Results were considered significant at the α = 0.05 level. Prior to hypothesis testing, the normality of dependent variables within each group was assessed using the Shapiro-Wilk test. Following verification of normality for every independent group, parametric tests were performed; otherwise, non-parametric analyses were performed. Parametric tests were conducted using a one-way ANOVA with post-hoc t-tests, while nonparametric tests used the Kruskal-Wallis test followed by Wilcoxon Rank Sum post-hoc analyses. Significance levels were adjusted for multiple comparisons using a Bonferroni correction.

### Volume Conductor Computational Model

A finite element method (FEM) computational model was developed to illustrate activation of on-target fibers (trigeminal nerve branches) versus off-target fibers (cutaneous nociceptors) (Verma, 2021a). The activating function, a proxy for nerve fiber engagement (Rattay, 1986), was used to characterize how the spatial distribution of the extracellular field from TENS stimulation influences activation of unmyelinated nociceptor fibers that run parallel to the skin surface, as well as to illustrate nerve fiber engagement within the SON, ION, and MN nerve trunks.

A quasistatic electric currents model was created in COMSOL 6.2 using the AC/DC Electric Currents (ec) module to solve for the electric potential within a three-layered homogeneous volume conductor under bipolar stimulation. The conductivity and permittivity values for each of the three layers were taken from a steady-state, non-capacitive model from (Verma, 2021a), which draws on low-frequency impedance data by Gabriel (1996). A highly conductive copper electrode (conductivity: 5.99 × 10⁷ S/m; permittivity: 1) with uniform current density was used to represent the TENS voltage source. Relative conductivity and permittivity values are provided in **Supplementary Table 4**.

It is well documented that the electrical properties of the skin are highly influenced by skin preparation (Tronstad, 2010; Verma, 2021a). We varied physical characteristics of the stratum corneum, the outer layer of skin, to understand how variance in the skin layer may impact on- vs off-target activation. A lumped stratum corneum impedance layer was modeled between the TENS contacts and the skin, with a thickness of 20 µm (Czekalla, 2019) and a permittivity equal to that of the skin. A sensitivity analysis of stratum corneum impedance was performed, set at one and two orders of magnitude greater and less than that of the skin, as well as its thickness, which ranged from 10 to 30 µm in 5 µm increments. This lumped layer was intended to simulate the effects of increased impedance due to layers of dead skin and sweat glands increasing local electrolyte concentration within the stratum corneum (Tronstad, 2010). Since noninvasive stimulation involves variable electrode conformance and limited compliance to deliver the necessary current, voltage-controlled stimulation was utilized in the model (Trout, 2023). Bipolar voltage sources of ±15 V were applied to the upper surfaces of the TENS copper discs. The lateral edges of the model, spanning the skin, adipose, and muscle layers, were assigned as ground, while all other external surfaces, including the skull, were treated with no-flux boundary conditions to simulate high-impedance barriers (Gabriel, 1996). Current conservation boundary conditions were applied throughout the continuous TENS-tissue domains.

Within the skin domain, a line 100 µm beneath and parallel to the skin surface was used to evaluate the activating function at a relevant nociceptor depth (Verdugo, 2022). Similarly, the activating function along the x-direction for the nerve trunk was calculated along the relevant nerve trunk depths for the trigeminal nerve branches positioned between the muscle and adipose layers. The activation function was used to illustrate the propensity of depolarization for nerve fibers at each of these depths within the tissue volume. Models were meshed using extremely fine free triangular meshes with a dynamic number of nodes distributed along the nociceptor and nerve evaluation lines. The number of elements was increased until the calculated activation function maxima fell within 1% of the value of the finest mesh. The thicknesses of each of these tissue layers, derived from average measurements in the cadaver cohort, can be found in **Supplementary Table 2**. Simulations were run on a Windows 10 computer with a 3.0 GHz Intel i7 processor and 32 GB of RAM.

## Results

We characterized nerve depth at the foramen, fascicle size, and fascicular to connective tissue ratios for each potential trigeminal nerve target: the SON, ION, and MN. Biophysics of electrical stimulation demonstrate that depth is a driving factor in nerve activation—fiber activation falls off with the square of the distance for bipolar stimulation (Plonsey and Barr, 2007). Additionally, increasing fascicle size increases activation thresholds (Davis, 2023; Grinberg, 2008) due to the increased resistivity from correlated perineurial thicknesses. From each of the five donors, we analyzed nerve samples from the right and left SON, ION, and MN. The right SON of D4 and the right MN of D5 were excluded due to previous traumatic injury. Qualitative anatomical details, including branching patterns for each donor, are described further below, and quantitative depths, average fascicular makeup, and areas are provided in **Supplemental Tables 3 and 4**.

### Supraorbital Nerve Anatomy

The SON exits the skull via the supraorbital foramen. Qualitative images of branching patterns after exiting the foramen are shown in **Figure 2A**. While the depths from the foramina are reported as fixed anatomical landmarks, the course of the trigeminal nerve afferents within the scalp after exiting the foramen will likely be engaged first by electrode stimulation. Additionally, the supraorbital foramen had differing amounts of bone between the foramen and the eye socket: some donors had a notch with no bony tissue between the nerve and the outer surface of the eye socket, and some had a true foramen. The average SON foramen to midline distance was 26.40±3.60 mm. Amongst the potential trigeminal nerve targets, the SON was the smallest, with an average circumference of 6.33±1.94 mm.

**Figure 2.**
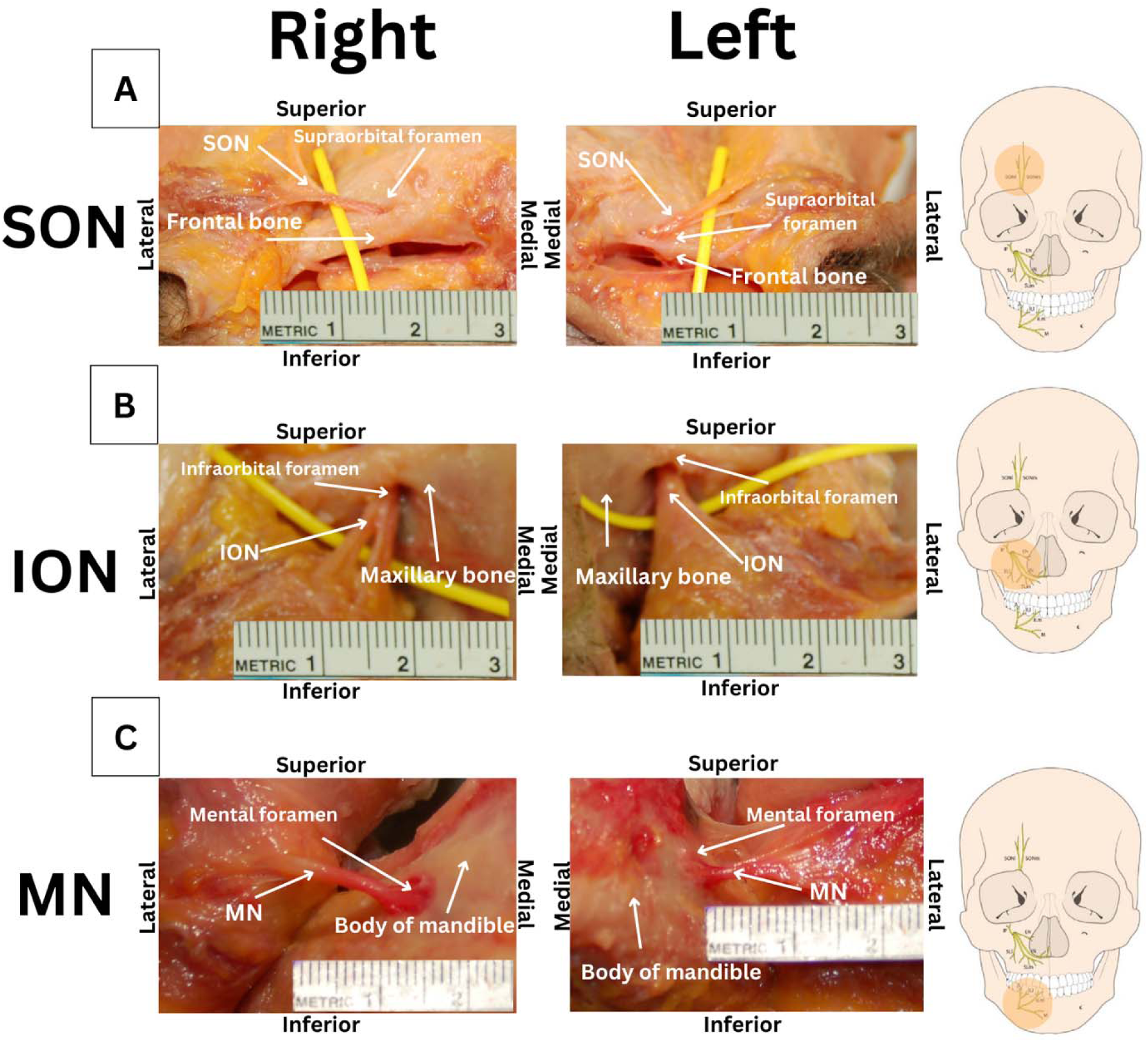
Dissection images from a single donor of the SON (A), ION (B), and MN (C) with accompanying skull diagrams indicating the trigeminal nerve branch of interest. Panel A shows the SON exiting the supraorbital foramen. These images show gross anatomical variations of the supraorbital foramen, with the right side having a greater thickness of bone between the foramen and the superior ridge of the eye socket. The SON begins to web and branch approximately 1 cm after exiting the foramen. Panel B shows the ION exiting the infraorbital foramen. These images also show ION anatomical variations on the right side immediately branching into two main branches, and the left side remaining in a single bundle. Panel C shows the MN exiting the mental foramen. Both sides were grossly similar in branching appearance. Corresponding skull diagrams were obtained with licensing permissions from https://creativecommons.org/licenses/by-nc/3.0/ with our addition of orange highlighted circles indicating the nerve branch of interest.

### Infraorbital Nerve Anatomy

As seen in **Figure 2B**, the right ION can be seen splitting almost immediately after exiting the skull, as opposed to the left side which remains as one nerve bundle contained in connective tissue. These gross anatomical differences varied bilaterally on the same donor as well as between donors. The ION had the greatest foramen to midline distance and had the most variation between donors, with an average distance of 27.52±5.45 mm. Of the three nerves, the ION had the second largest average circumference at 17.81±9.61 mm.

### Mental Nerve Anatomy

The MN exits the skull via the mental foramen in the body of the mandible and starts to branch out after exiting the foramen (**Figure 2C**). The MN had an average foramen to midline distance of 21.92±2.60 mm, and the average circumference was 19.46±7.87 mm. No remarkable hemispheric differences in branching pattern or foramen anatomy were noted in our donors.

### Comparative Anatomy of Trigeminal Nerve Branches, Nerve Depths, and Fascicular Composition Across Branches

Here, we were interested in comparing the parameters most relevant for transcutaneous stimulation: nerve depth and fascicular size (Grinberg, 2008). The SON was, on average, the most superficial nerve, with the depth from the skin to the foramen being 6.15±0.90 mm, and it had the least variation in depth between donors. The ION was deeper than the SON with an average depth of 9.65±1.43 mm. The MN had an average depth of 9.85±2.87 mm and had the greatest variance in depth among donors. Depth for the MN was measured from the surface of the skin, not the mucosal surface, to the foramen during dissection. ANOVA found a significant difference in branch depth when comparing the SON, ION, and MN (p = 0.0001). Pairwise post hoc t-tests revealed that the SON courses more superficially than both the ION (p = 0.00068) and the MN (p = 0.00036), positioning it as the best of the three candidate branches for transcutaneous targeting.

The SON had the smallest number of fascicles, with an average of 5.22±2.11 fascicles per nerve sample, and the average individual fascicular area was 0.0997±0.100 mm^2^. Across nerves, Kruskal-Wallis revealed no significant difference between individual fascicle areas (p=0.06945).

While the SON had the smallest number of fascicles, it had the largest fascicular tissue to connective tissue ratio, as shown in **Figure 3A**. The ION had the greatest number of fascicles, averaging 16.1±6.67 fascicles and an average individual fascicular area of 0.0931±0.1026 mm^2^. The MN had an average of 13±5.12 fascicles per nerve sample. It also had the lowest ratio of fascicular tissue to connective tissue, as displayed in **Figure 3C**, and it had the lowest average individual fascicular area at 0.0855±0.1062 mm^2^.

**Figure 3.**
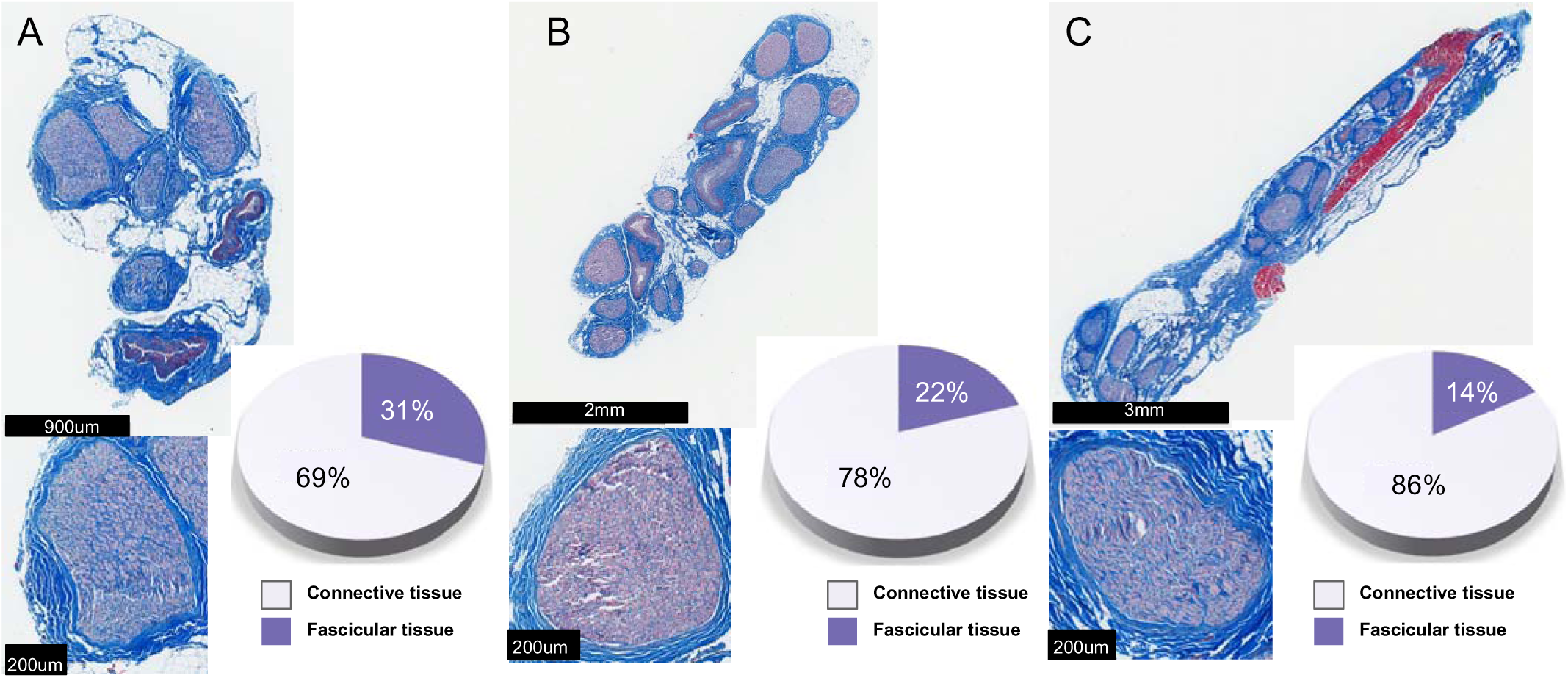
Histology of the SON (A), the ION (B), and the MN (C), with accompanying close-up images of a representative fascicle from each nerve below, and the average percentage amounts of connective tissue to fascicular tissue for each nerve. The SON had an average fascicular tissue makeup of 31.4%±10.0%, followed by the ION and the MN.

The proportion of fascicular tissue to connective tissue was measured for each nerve in the five human donors, and average values were compared across the cohort. Connective tissue is a high impedance pathway for stimulation and thus may influence activation of neural tissue. The SON had the lowest proportion of connective tissue, with an average makeup of 31.4±10.0% fascicular tissue. The ION had the next largest ratio with 22.4±11.7% fascicular tissue, while MN had the smallest ratio with 13.5±7.35% fascicular tissue. ANOVA revealed a significant difference in fascicular area breakdown when comparing the SON, ION, and MN (p=0.0316). Multiple comparisons showed pairwise differences between SON and MN (p=0.0023), but not between the SON and ION (p=0.1814), nor between the MN and ION (p=0.1845), indicating that the SON contains more fascicular area by percent makeup than the MN.

### Volume Conductor Nerve Activation Modeling Results

To illustrate activation of nerve fibers within trigeminal nerve trunks at the depths consistent with our anatomical findings, we created three FEM models of the associated SON, ION, and MN tissue layers (**Figures 4A-C**). **Figure 4D** shows a representative model of the activation function along the x-direction in the SON. Maximum values of the activation function along the x-direction for nociceptors within the skin and nerve trunks were evaluated for each of the three models shown in **Figure 4A-C**. Regions of positive activation function (red) illustrate regions of increasing propensity for depolarization or activation. Overlaid vector fields represent current density, and black circular arcs near the TENS contacts denote equipotential zones, illustrating the spread of the electric field. The activation function magnitude decreases steeply with distance from the TENS contacts, showing orders of magnitude variation between where superficial nociceptive fibers are found compared to fibers located deeper, within the trigeminal nerve trunks (**Figure 4D**).

**Figure 4.**
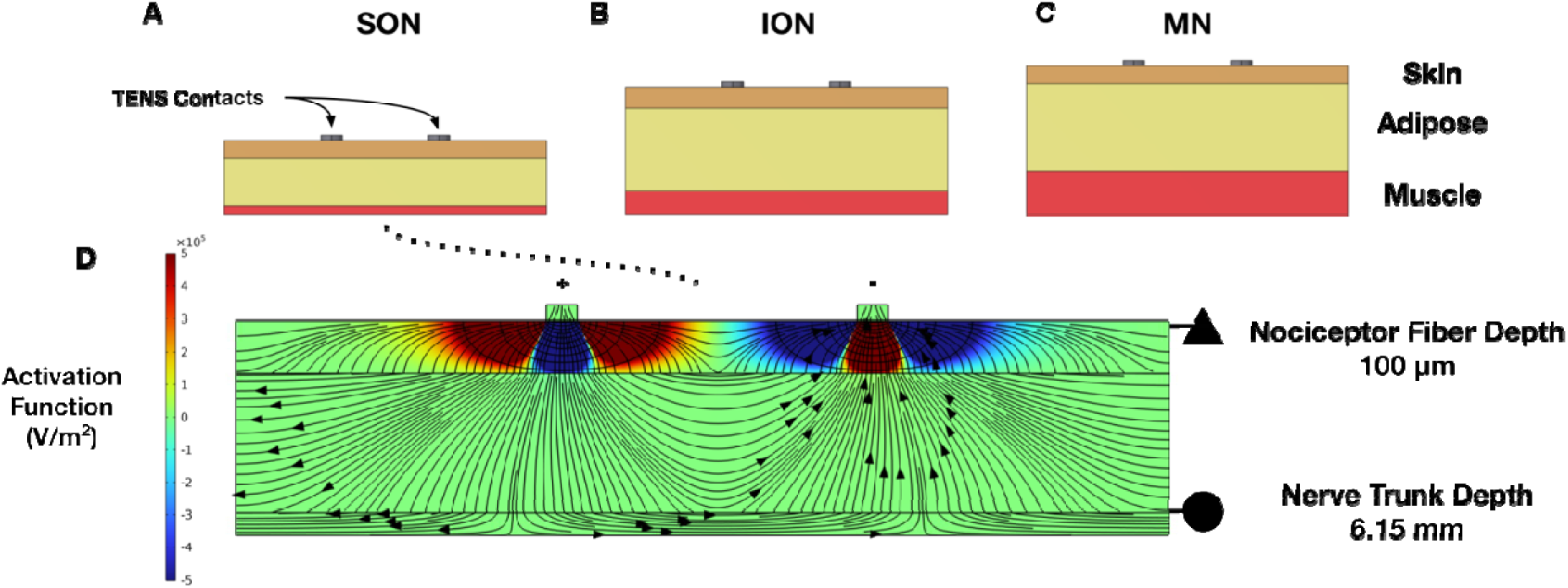
Computational Modeling of the Activation Function during Bipolar Transcutaneou Electrical Nerve Stimulation (TENS) of Trigeminal Nerve Branches. (A–C) Average anatomically derived depths of the supraorbital nerve (SON), infraorbital nerve (ION), and mental nerve (MN) tissue, respectively. (D) Simulated activation function (color gradient, red: activation) in the SON across a tissue cross-section during bipolar TENS for each of the three branches, shown along the x-direction. The color scale in Figure 4D is capped at 5×10⁵ for visualization; peak values are detailed in Figure 5.

### Impact of Skin Thickness: On vs. Off-Target Activation Function Magnitude

Because skin composition and preparation have a drastic effect on electrode-skin interface properties (Tronstad, 2010; Verma, 2021a), we modulated properties of the stratum corneum-TENS contact interface, thickness and impedance for each of these three computational models, and compared the maxima of the activation function at the level of nociceptive fiber and nerve trunks. Despite variance in stratum corneum impedance and thickness, depth wa the largest driver of the maximum activation function (level of circles), seen in Figure 5D/E-zoomed panel. (SON being significantly closer to the surface as compared to the ION and the MN). Nociceptive activation function values were orders of magnitude greater than those of nerve trunk activation function values. While the activation function is orders of magnitude different between nociceptive fibers and nerve trunk fibers, the activation function magnitude necessary to activate smaller diameter fibers associated with nociceptive Aδ and C fibers i typically higher (canonically 5-10x as a function of the difference in interaxonal diameter) (Warman, 1992; Danner, 2011; Yoo, 2013; Nicolai, 2020). In the nociceptor zone, increasing stratum corneum barrier thickness decreased the activation function linearly (Figure 5D/E-triangles), while deep nerve trunks were less affected (Figure 5D/E-zoomed circles). Increasing stratum corneum impedance preferentially decreased activation of nociceptors. Distance engagement remains a driving factor in this illustrative model of anatomically realistic tissue depths for on- vs-off target engagement.

**Figure 5.**
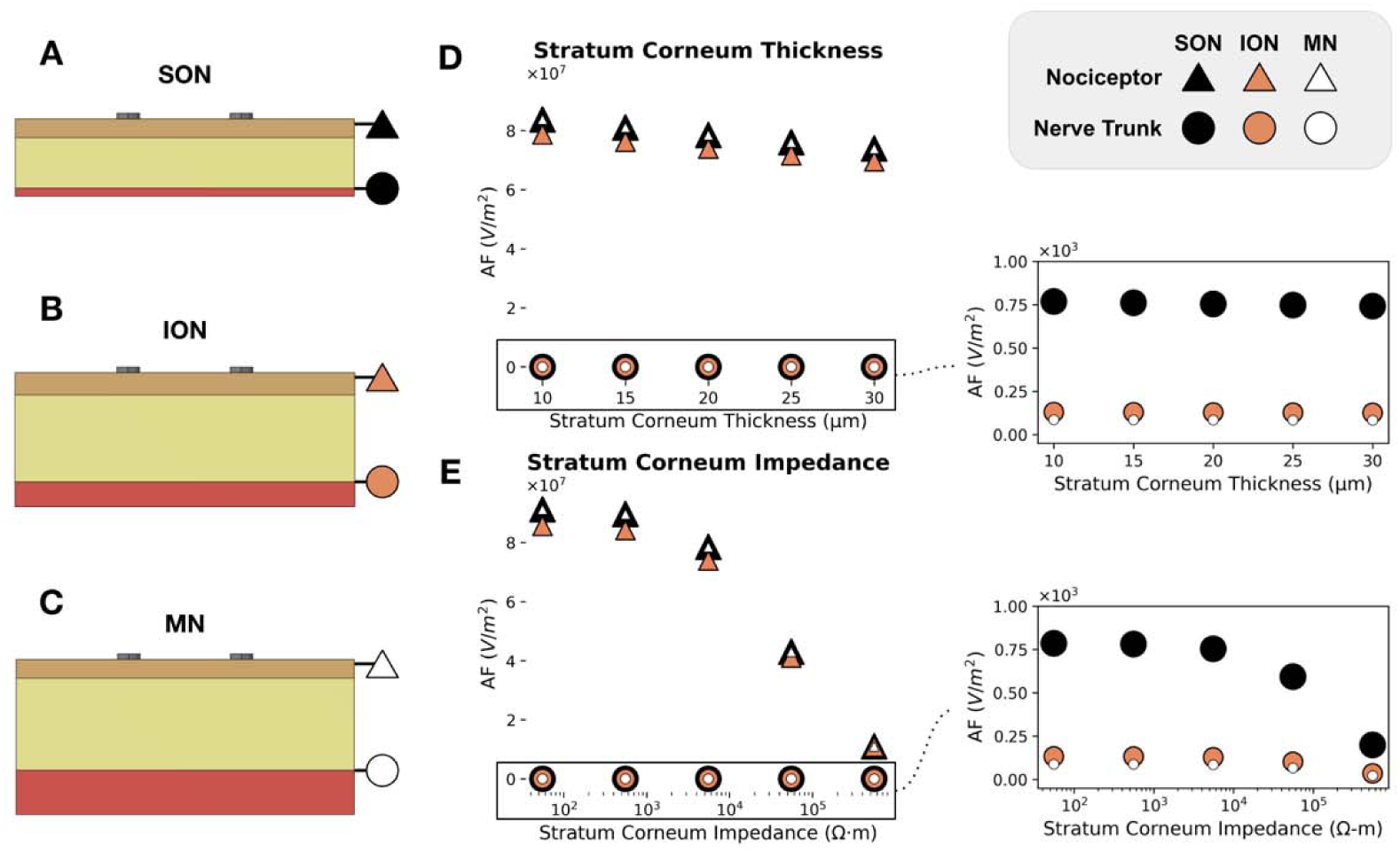
Influence of Stratum Corneum Properties on Nociceptor and Trigeminal Nerve Trunk Activation during TENS Sensitivity Analysis. (A–C) Schematic representations of the supraorbital (SON), infraorbital (ION), and mental (MN) nerve branches beneath the skin, illustrating the position of superficial nociceptors (triangle) and deeper nerve trunks (circle). (D) Effect of stratum corneum thickness (10–30 µm) on peak activating function (AF) values at nociceptor and nerve trunk depths. (E) Effect of stratum corneum lumped barrier impedance (1 and 2 orders of magnitude larger and smaller than that of the scalp) on peak AF values. Parameterization of the stratum corneum, modeled as a thin lumped impedance layer between the TENS contacts (grey) and scalp (tan), significantly modulates activation function magnitude, preferentially for nociceptors, and secondarily for nerve trunk depth.

## Discussion

The trigeminal nerve is a promising cranial nerve neuromodulation target for modulating autonomic tone and cerebral blood flow, with several favorable features for noninvasive neuromodulation. Stimulation of the trigeminal nerve has elicited many documented effects on blood flow changes (Chiluwal, 2017), metabolism, and inflammation (Powell, 2023); however, stimulation parameters and anatomical characteristics are variable, and therapeutic criteria for engagement are varied across doses, timing, electrode placement, and efficacy (Shiozawa, 2014; Powell, 2023; Westwood, 2023). As a transcutaneous target for neuromodulation, little is known about the human trigeminal nerve fascicular composition, gross anatomical landmarks such as the locations of its foramina, and anatomical depths of its trunks. These factors are important for neuromodulation as these nerve targets exit and branch into cutaneous end-organ targets in the face. This study summarizes anatomical measurements of the trigeminal nerve, including CT-derived anatomical data regarding foramen midline distances, microdissection nerve depths at defined foraminal landmarks, fascicular tissue makeup, and an illustrative model of nerve activation at the level of nociceptors and average nerve trunk depths. From these results, the SON emerges as a favorable transcutaneous neuromodulation target of the three trigeminal nerve branches examined, due to its high nerve to connective tissue composition at the foramen relative to the ION and MN, its sensory only nature, and superficial course (∼6mm) relative to the cervical vagus (Hammer, 2018). These findings are consistent with small animal, large animal, and human studies, wherein stimulation of the SON creates the most consistent/repeatable changes in blood flow, with the MN having the least consistent data across models. (White, 2021)

### Comparative advantages of the SON to the ION and MN

#### SON Depth Influences Nerve Trunk Activation

The trigeminal nerve meets several criteria as a favorable neuromodulation target for non-invasive stimulation of the nerve trunk. On average across the cohort, the SON trunk runs 6.15mm below the surface of the scalp at the foramen. We found that the supraorbital nerve follows a branching path ∼1cm after exiting the foramen above the eye socket in the cranium and is statistically more superficial than the ION and MN branches (p<0.005). This is in contrast to the depth of the cervical vagus reported in the literature, which is approximately 36cm from the surface of the skin (Hammer, 2018). Superficial nerves, such as the SON, can have better TENS engagement than the ION and MN simply due to their location, barring local density of nociceptors (Morch, 2011). Furthermore, the SON contains purely sensory fibers, in theory making it easier to selectively activate on-target, Aβ/Aδ sensory afferents, as low-threshold, off-target motor efferents are localized within the facial nerve. The location of the SON is also favorable from an aesthetic standpoint. Electrodes placed on the brow are more easily concealed with a hat or scarf than electrodes on the cheek or chin (DeGiorgio, 2006). For animal cortical validation and accessibility of the SON in invasive neuromodulation, previous studies of the rat SON and ION show variations in nerve size and accessibility underneath soft tissue between rats and mice (Dingle, 2019). We emphasize careful consideration of relative current thresholds when evaluating these nerves as transcutaneous nerve targets in animal models for human translatability.

#### Fascicular Tissue Histology

While we did not differentiate types of tissue within the umbrella of connective tissue, high impedance perineurium and connective tissue layers require higher stimulation currents for activation at the same depth, which could increase the likelihood of unwanted off-target effects (Grinberg, 2008). Fascicular size also plays a role in activation thresholds, with smaller diameter fascicles having a lower activation threshold compared to larger diameter fascicles (Grinberg, 2008; Davis, 2023). Additionally, in animal models of the ION, there was significant variability in extracellular matrix (ECM) and nerve tissue between rats and mice (Dingle, 2019), indicating that caution should be taken in creating anatomically realistic models for determining activation thresholds for specific species. In our histological analysis, the SON was found to have a significantly higher ratio of fascicular tissue to connective tissue when compared to the MN (p =0.0023), but not when compared to the ION (p=0.1814). Across the cohort, the SON had approximately 31.4% average fascicular area makeup, the largest of the three branches examined.

### Computational model of nerve trunk engagement for neuromodulation

From our anatomical measurements, we found the SON trunk had a significantly more superficial course compared to the other trigeminal branches at the foramen. This is an important consideration for transcutaneous stimulation, as the depth of the nerve trunk significantly influences activation of the nerve fibers. Using the activation function (the second-order spatial derivative of the extracellular potential) as a proxy for fiber engagement (positive: depolarization) (Rattay, 1986), we found that activation function where superficial nociceptor associated fibers lie (∼100 µm depth) was orders of magnitude higher than that at deeper nerve trunks (6.15–9.85 mm), which lie between the adipose and muscle layer. We varied electrode size, spacing, and stratum corneum conductivity and found that increased stratum corneum impedance and thickness (i.e., due to skin hydration variability or dead skin) reduced nociceptor activation while only minimally affecting deeper nerve engagement. Of the nerve branches examined, the SON was most susceptible to changes in activation when varying lumped parameters of the stratum corneum, but not as significantly as nociceptor variance. Sweating, inconsistency in skin hydration, and electrode-skin compliance session to session can impact characteristics of the stratum corneum (Czekalla, 2019; Lu, 2018).

### Comparing cranial nerve stimulation modalities

TNS presents a promising target for transcutaneous neuromodulation of cranial nerves due to the superficial course of its SON, ION, and MN branches, as well as its primarily sensory composition. However, its clinical utility is not yet optimized due to a lack of anatomical localization of these branches and a lack of objective stimulation parameters. Off-target effects, including unwanted stimulation of neighboring nociceptors, can be treatment-limiting and result in less effective nerve stimulation, as seen with other cranial nerve neuromodulation modalities. For example, invasive vagus nerve stimulation (iVNS) is one of the most extensively studied and clinically validated methods. However, risks of iVNS, including surgical complications and voice changes, make this procedure not suitable for all patients (Ben-Menachem, 2001). Additionally, since the vagus nerve is a mixed motor and sensory fiber nerve, low-threshold Aα motor fibers have been implicated in therapy-limiting off-target effects, including motor activation of neck muscles, dyspnea, cough, and arrhythmias (DeFerrari, 2017; Zannad, 2015; Nicolai, 2020; Settell, 2020). Noninvasive transcutaneous vagus nerve stimulation has been utilized at the superficial auricular branches of the vagus near the ear (taVNS), but with variable reported outcomes in clinical applications (Yap, 2020; Verma, 2021b). taVNS targets sensory afferent fibers from the auricular branch of the vagus nerve as they innervate the skin, but anatomical studies show that this branch loses fibers to facial nerve cross-connections and forms a dispersed plexus, rather than a trunk near the ear (Verma, 2021b). As a result, taVNS likely stimulates only a small subset of vagal axons located directly beneath the electrode and not the nerve trunk itself (Verma, 2021b). Similarly, transcutaneous stimulation of the cervical vagus nerve (tcVNS) has been studied clinically, but selective activation is likewise limited by off-target effects from nearby nociceptor and motor fiber activation (Yap, 2020) including potential cardiac effects due to inadvertent activation of efferent fibers to the heart when the vagus is stimulated bilaterally (Colzato and Vonck, 2017).

### Cranial Nerves Share Projections and Nuclei for Processing and Ameliorating Neurological Conditions

Both tVNS and TNS have been utilized clinically in the treatment of depression and drug-resistant epilepsy and may share a common bottom-up mechanism where activation of brainstem nuclei causes long-term modifications of cortical circuitry (Colzato and Vonck, 2017). While the exact mechanisms of central processing are under debate (Zhang, 2020; Cook, 2013; Schrader, 2011; Suminski, 2018), the projections from the vagus afferent pathways share nuclei with those of the trigeminal nerve.

The vagus and trigeminal nerves share connectivity to the nucleus tractus solitarius (NTS) (Huff, Weisbrod and Daly, 2025) (Patel, Jozsa and Das, 2025), which is linked to processing visceral sensory information, as well as the ventroposteromedial nucleus (VPM) (Mithani, 2020). Neuromodulation aims to tap into these pathways as a form of biomimicry. Activation of the SON and ION can also leverage afferent input to the SPG to activate the dive reflex, which modulates cerebral blood flow and CSF penetration as a neuroprotective effect (Lim, 2023). As opposed to the vagus nerve, the SON and ION branches are purely sensory in composition (Powell, *et al*., 2023), containing Aβ and Aδ fibers innervating the face, and have superficial exit points from the skull (Netter, 2019). In TNS, depolarization of fibers within the trunk does not directly innervate off-target muscles implicated in therapy-limiting side effects (Nicolai, 2020); rather, motor side effects are attributed to activation of nearby nerves innervating facial muscles and nociceptive sensation from activation of cutaneous pain fibers.

Despite these commonalities in projections, the natural sensory traffic from the vagus nerve encodes sensory information from the neck muscles and all the visceral organs. In contrast, the natural sensory traffic from the trigeminal nerve encodes sensory information from the skin and muscles of the scalp, forehead, and face that includes proprioceptive information, temperature, and wetness, (Huff, 2025) which may influence the dive reflex. Consequently, it should not be presumed that because the vagus and trigeminal nerves have similar connectivity within the brain that their responses within the brain are identical. Unlike the trigeminal nerve, the vagus nerve also includes efferent outflow to the neck muscles as well as parasympathetic outflow to most of the visceral organs. Due to its much greater diversity of function, isolating only the afferent traffic to the brain may be much more difficult for VNS. This was shown in several recent failed clinical trials, where therapeutic doses were not achieved due to limiting off-target effects (DeFerrari, 2017; Zannad, 2015; Nicolai, 2020; Settell, 2020). Additionally, activation of the sympathetic trunk, a hitchhiking nerve adjacent to the vagus, likely leads to further off-target activation for some indications (Deshmukh, 2025). This motivates a renewed interest in alternative cranial nerve pathways with similar brain connectivity but potentially fewer sources of limiting off-target activation.

## Limitations

### Intersubject Variability

While we make generalizable statements about the course of the trigeminal nerve, there was anatomical variability between donors, and our donor pool has limitations. All donors used were male; future studies can look at female donors and the anatomical differences between sexes. One donor had a previous jaw fracture, and another had previous eye socket trauma, resulting in missing measurements for those nerves and foramina, and potentially increasing variance across the cohort for measurements of nerve depth. Additionally, some donors had a supraorbital notch as opposed to a supraorbital foramen. While it is unclear if this would affect activation of the nerves, it is a variable to consider when placing electrodes and determining the distance from midline. The ION and MN had the greatest variability of nerve depth, which could be due to the variance of fat distribution in the face; below the eye and on the chin typically have more adipose tissue than above the eye, on the brow (Donofrio, 2000). The average BMI of the cadavers was 25.388±2.952. According to the National Health and Nutrition Examination Survey (NHANES), the average BMI of Americans was 29.23 in 2023 (Young, 2023). This may be a limitation to our measurements for generalizability in trigeminal nerve targeting towards patients in the general population, since the adipose tissue between the scalp and the nerve trunk was based on average nerve depths from our cohort, which is also utilized in the computational modeling component of the study regarding the activation function.

### Imaging and Histological Limitations

All of our measurements are based on internal landmarks and CT scans. External landmarks will be relevant for electrode placement in a clinical setting to avoid CT or MR imaging which could increase associated patient costs. Future studies should include external landmarks to ensure precise transcutaneous electrode placement to increase clinical applicability and provide a more practical map of the trigeminal nerve. While it is possible to palpate the supraorbital, infraorbital, and mental foramina, additional factors must be considered when placing electrodes in terms of location and physical properties of the skin.

Fascicular area and the ratio of fascicular tissue to connective tissue measurements were taken from nerve samples immediately after the nerves exited the skull through their respective foramina. Due to the elasticity of the nerves, the exact point where the nerve exits the foramen where the tissue sample was harvested may have some variability. More importantly, the sample was taken proximal to where the nerves begin branching and distributing across the face. Because of this, it is unknown how the ratio of fascicular tissue to connective tissue changes as we move distally along the nerve branches. Additionally, the rotation of certain fascicles closest to the surface of the skin at the foramen does not necessarily indicate the most superficial and hence most easily depolarized fascicles.

### Computational Model Limitations

While we can infer relative depolarization for unmyelinated fibers distributed arbitrarily within a volume of tissue, there are several limitations regarding our biophysical model. The model is a perfectly conductive and homogenous medium. Modeling nerve fibers diving in and out of the muscle fascia could further refine the representation of nerve fibers that lie between muscle and adipose tissue layers with anisotropic current spread. Additionally, the exact depolarization thresholds of different fiber types will heavily depend on myelination, ion channel density, and capacitance of the membrane (Rattay, 1986; McIntyre, 2002). Exact activation thresholds must consider the frequency domain, as well as biophysical constants such as the membrane resistance, myelination, and most prominently, the orientation, branching, and distribution of nociceptors near each of these TENS contacts (Morch, 2011). Additionally, the model does not capture the difference in activation thresholds between Aβ and Aδ fibers, which arise from their differing diameters and internodal spacing (Plonsey and Barr, 2007). Future work could incorporate cable properties of known fiber types (Rattay, 1986; McIntyre, 2002) within the trigeminal branches and patient-specific anatomical features to better refine predictions of activation thresholds (Sweeney, 1990; Bucksot, 2021; Blanz, 2023; Musselman, 2023). Further, the activation function at nociceptive fiber-relative depths also does not capture perceptive pain thresholds. The model created can illustrate activation for a generic neural fiber within the context of the anatomical depths we described for each of the branches of the trigeminal nerve.

Despite clear trigeminal nerve courses, the major obstacles in understanding on- vs. off-target engagement of sensory fibers in the scalp are the locations of nearby facial nerves and muscles, and the density, relative distribution, and rotation of nociceptive fibers in evoked nociceptive sensation. Since we solely looked at the trigeminal nerve course, future studies should include the dissection and mapping of the nearby facial nerves and the efferent fibers that innervate facial muscles. Understanding the course of these nerves would minimize off-target muscle contractions and optimize electrode placement.

## Conclusion

This study assessed the anatomical basis of transcutaneous trigeminal nerve stimulation (TNS) by characterizing three accessible peripheral branches: the supraorbital nerve (SON), infraorbital nerve (ION), and mental nerve (MN). We provided quantitative measurements of nerve depth, fascicular area, connective tissue composition, and foramina-to-midline distances using microdissection and CT imaging. The SON was found to have the most superficial foramina among the branches examined. A finite element model was used to evaluate how anatomical differences influence nerve engagement, showing nerve trunk engagement is primarily driven by distance from the stimulation site, and that superficial fibers, particularly those in the SON, are most sensitive to changes in skin impedance properties such as stratum corneum thickness and impedance. The model’s sensitivity and large activation function for superficial nociceptors serve as a caution that off-target nociceptive activation can dominate in, and is likely dependent on, density and location, warranting further study. This study serves as a guide for further investigation of transcutaneous electrode development and computational modeling for targeted neuromodulation in clinical practice.

## Acknowledgements and Disclosures

Special thanks to Nishant Verma and James Trevathan for FEM modeling advice and to Imaging Services at the Wisconsin Institute for Medical Research for assistance with CT imaging of the donor tissue. This work was funded by the Defense Advanced Research Projects Agency (DARPA) Biological Technologies Office (BTO) Targeted Neuroplasticity Training Program under the auspices of Doug Weber and Tristan McClure-Begley through the Space and Naval Warfare Systems Command (SPAWAR) Systems Center with (SSC) Pacific grants no. <u>N66001-17-2-4010</u>.

KAL and JCW are co-founders and equity holders of NeuraWorx, a privately held neuromodulation company investigating cranial nerve stimulation for glymphatic clearance. KAL is also a co-founder and equity holder for Neuronoff, Inc. KAL is a scientific board member and has stock interests in NeuroOne Medical Inc. KAL is also a paid member of the scientific advisory board of Abbott and Presidio Medical, and a paid consultant for the Alfred Mann Foundation, ONWARD and Restora Medical.

## Supplementary Materials

**Supplementary Table 1.** Tissue conductivity and permittivity parameters used in three-layer trigeminal nerve computational model

| Tissue Layer | Conductance (S/m) | Permittivity |
| --- | --- | --- |
| Skin | 1.8e-4 | 1.17e3 |
| Adipose | 2.46e-2 | 2.08e4 |
| Muscle | 5.23e-1 | 1.24e6 |

**Supplementary Table 2.** Average tissue depths used in COMSOL FEM models

| Nerve | Scalp thickness (mm) | Adipose thickness (mm) | Corresponding muscle | Muscle thickness (mm) | Nociceptor depth (mm) | Nerve depth (mm) |
| --- | --- | --- | --- | --- | --- | --- |
| SON | 1.67 | 4.48 | Frontalis | 0.73 | 0.10 | 6.15 |
| ION | 1.97 | 7.68 | Orbicularis Oculi | 2.20 | 0.10 | 9.65 |
| MN | 1.67 | 8.18 | Mentalis | 4.00 | 0.10 | 9.85 |

**Supplementary Table 3.** Anatomical measurements of trigeminal nerve branches that were taken during microdissections of five human donors (D1-5). Measurements were not obtained from the right SON of D4 and right MN of D5 due to previous trauma to the respective areas. The bottom row shows the averages for each nerve across donors.

|  | Nerve depth (mm) |  |  |  |  |  | Distance from midline (mm) |  |  |  |  |  | Nerve circumference (mm) |  |  |  |  |  |
| --- | --- | --- | --- | --- | --- | --- | --- | --- | --- | --- | --- | --- | --- | --- | --- | --- | --- | --- |
|  | SON L | SON R | ION L | ION R | MN L | MN R | SON L | SON R | ION L | ION R | MN L | MN R | SON L | SON R | ION L | ION R | MN L | MN R |
| D1 | 5 | 5 | 7.5 | 7 | 14 | 14 | 29.1 | 32.7 | 26.9 | 33.9 | 20.1 | 22.8 | 7.86 | 9.87 | 38.9 | 15.6 | 30.8 | 6.57 |
| D2 | 7 | 6 | 11 | 11 | 7 | 6 | 28.0 | 22.9 | 21.4 | 36.1 | 21.4 | 25.1 | 3.63 | 4.86 | 22.9 | 11.4 | 23.8 | 29.3 |
| D3 | 7 | 7.5 | 9.5 | 9 | 10 | 10 | 23.4 | 26.3 | 24.2 | 22.7 | 17.1 | 21.3 | 4.24 | 6.78 | 9.79 | 9.93 | 15.6 | 14.6 |
| D4 | 7 | 6 | 10.5 | 11 | 9 | 7 | 27.9 | -- | 21.7 | 34.2 | 21.0 | 23.1 | 5.86 | -- | 11.3 | 10.0 | 22.7 | 18.3 |
| D5 | 6 | 5 | 10 | 10 | 9 | 12.5 | 20.9 | 26.3 | 26.3 | 27.9 | 25.6 | -- | 6.59 | 7.31 | 24.4 | 24.1 | 13.5 | -- |
| Avg | 6.4 | 5.9 | 9.7 | 9.6 | 9.8 | 9.9 | 25.9 | 27.1 | 24.1 | 31.0 | 21.0 | 23.1 | 5.63 | 7.20 | 21.44 | 14.18 | 21.3 | 17.20 |

**Supplementary Table 4.**
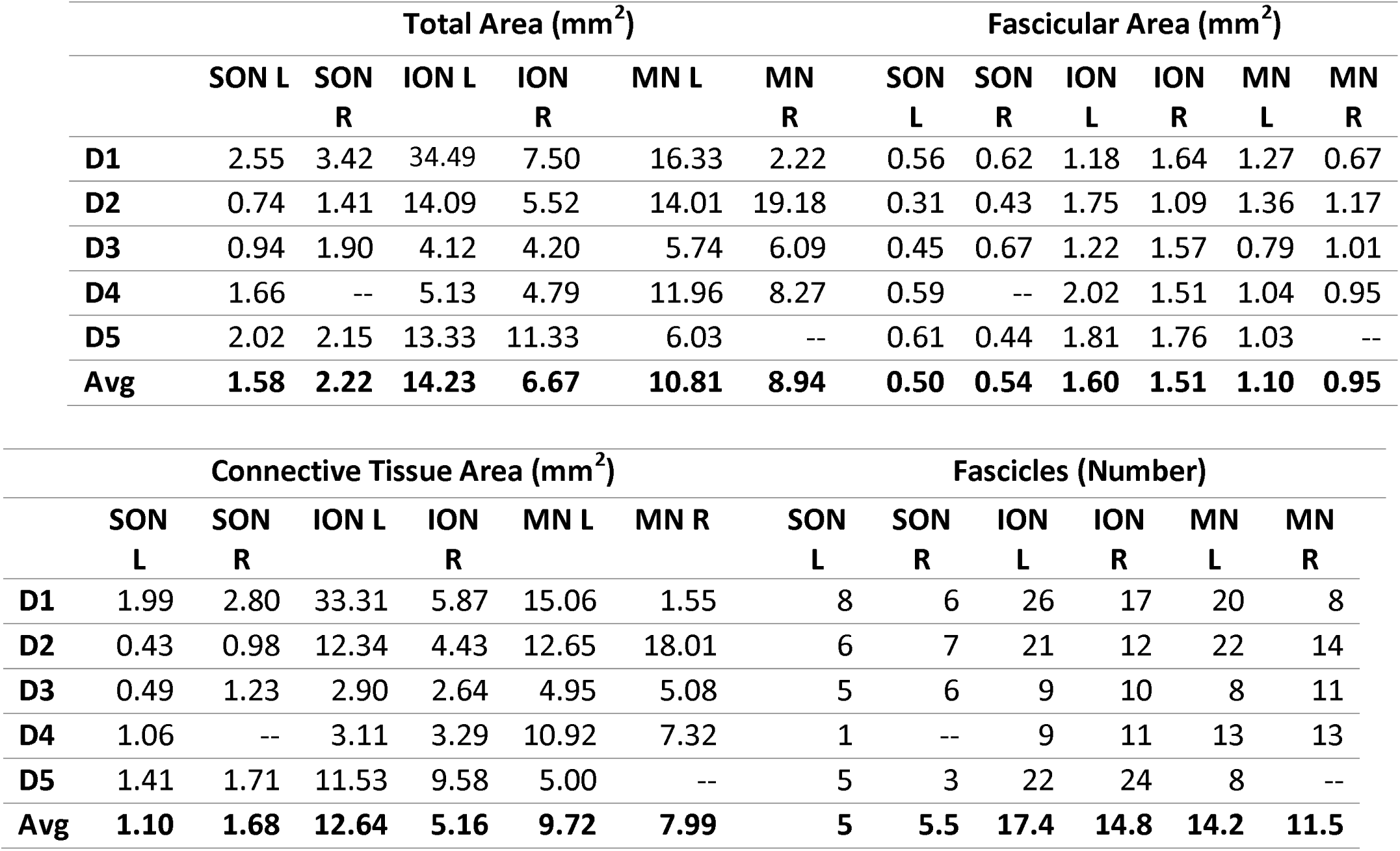
Histological measurements of trigeminal nerve sections at the foramen from five donors. Histological slices were taken from the SON, ION, and MN approximately where the nerve exited from each respective foramen. Total area, in millimeters squared, represents the area of each nerve sample for both left and right sides. The fascicular area, in millimeters squared, represents the total fascicular tissue area of all fascicles in each nerve sample. Connective tissue area, in millimeters squared, represents the fascicular area subtracted from the total area and includes everything in each nerve sample that is not fascicular tissue. Measurements were not able to be obtained for the right SON of D4 and right MN of D5 due to previous trauma. The bottom row shows the averages for each nerve across donors.

|  | Total Area (mm<sup>2</sup>) |  |  |  |  |  | Fascicular Area (mm<sup>2</sup>) |  |  |  |  |  |
| --- | --- | --- | --- | --- | --- | --- | --- | --- | --- | --- | --- | --- |
|  | SON L | SON R | ION L | ION R | MN L | MN R | SON L | SON R | ION L | ION R | MN L | MN R |
| **D1** | 2.55 | 3.42 | 34.49 | 7.50 | 16.33 | 2.22 | 0.56 | 0.62 | 1.18 | 1.64 | 1.27 | 0.67 |
| **D2** | 0.74 | 1.41 | 14.09 | 5.52 | 14.01 | 19.18 | 0.31 | 0.43 | 1.75 | 1.09 | 1.36 | 1.17 |
| **D3** | 0.94 | 1.90 | 4.12 | 4.20 | 5.74 | 6.09 | 0.45 | 0.67 | 1.22 | 1.57 | 0.79 | 1.01 |
| **D4** | 1.66 | -- | 5.13 | 4.79 | 11.96 | 8.27 | 0.59 | -- | 2.02 | 1.51 | 1.04 | 0.95 |
| **D5** | 2.02 | 2.15 | 13.33 | 11.33 | 6.03 | -- | 0.61 | 0.44 | 1.81 | 1.76 | 1.03 | -- |
| **Avg** | **1.58** | **2.22** | **14.23** | **6.67** | **10.81** | **8.94** | **0.50** | **0.54** | **1.60** | **1.51** | **1.10** | **0.95** |

|  | Connective Tissue Area (mm<sup>2</sup>) |  |  |  |  |  | Fascicles (Number) |  |  |  |  |  |
| --- | --- | --- | --- | --- | --- | --- | --- | --- | --- | --- | --- | --- |
|  | SON L | SON R | ION L | ION R | MN L | MN R | SON L | SON R | ION L | ION R | MN L | MN R |
| **D1** | 1.99 | 2.80 | 33.31 | 5.87 | 15.06 | 1.55 | 8 | 6 | 26 | 17 | 20 | 8 |
| **D2** | 0.43 | 0.98 | 12.34 | 4.43 | 12.65 | 18.01 | 6 | 7 | 21 | 12 | 22 | 14 |
| **D3** | 0.49 | 1.23 | 2.90 | 2.64 | 4.95 | 5.08 | 5 | 6 | 9 | 10 | 8 | 11 |
| **D4** | 1.06 | -- | 3.11 | 3.29 | 10.92 | 7.32 | 1 | -- | 9 | 11 | 13 | 13 |
| **D5** | 1.41 | 1.71 | 11.53 | 9.58 | 5.00 | -- | 5 | 3 | 22 | 24 | 8 | -- |
| **Avg** | **1.10** | **1.68** | **12.64** | **5.16** | **9.72** | **7.99** | **5** | **5.5** | **17.4** | **14.8** | **14.2** | **11.5** |

**Supplemental Figure 1.**
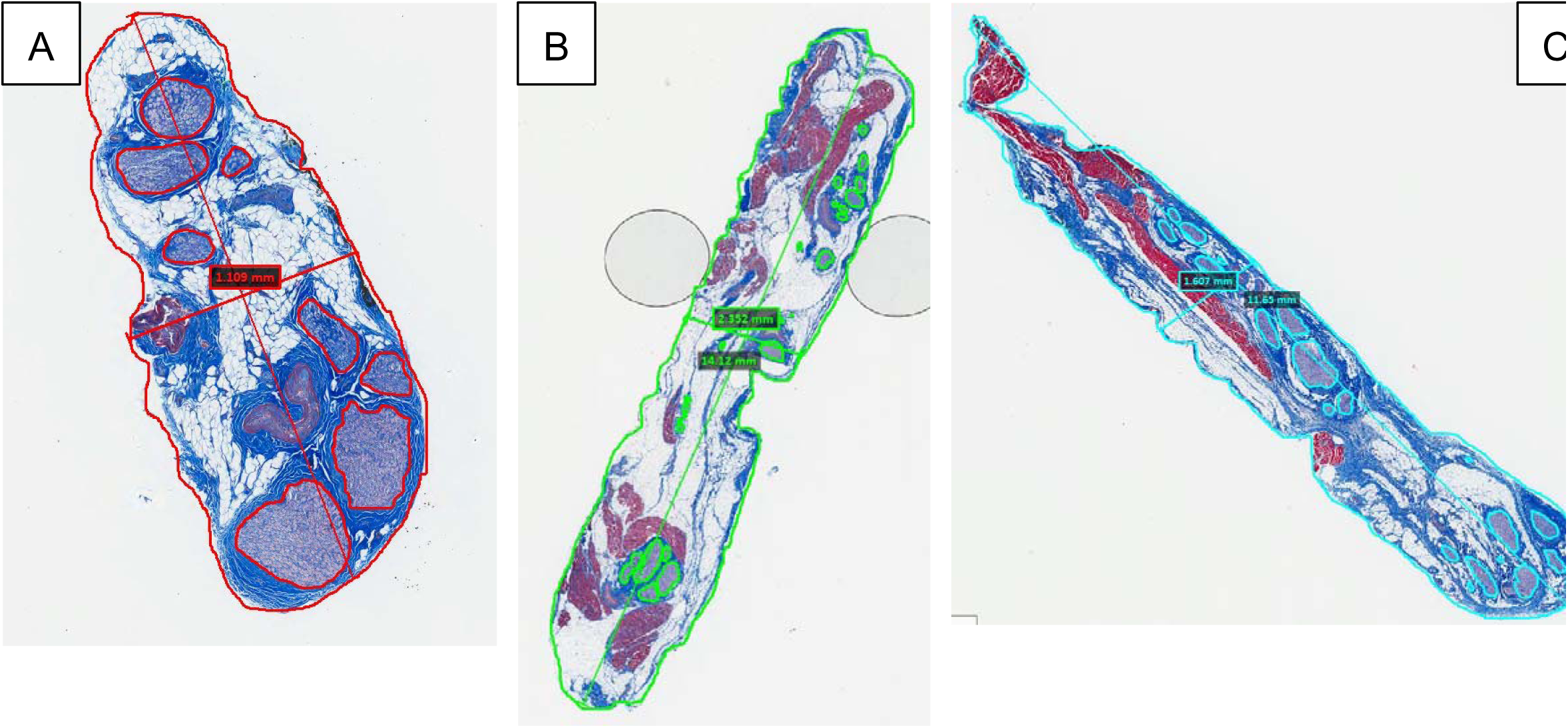
Examples of fully annotated nerve samples of the SON (A), ION (B), and MN (C). The entire circumference of the nerve was manually traced using the Aperio ImageScope drawing annotation tool. Each fascicle was then counted and manually traced. Lastly, the ruler tool was used to measure the length and width of each nerve.

